# Viscoelastic mapping of mouse brain tissue: relation to structure and age

**DOI:** 10.1101/2020.05.11.089144

**Authors:** Nelda Antonovaite, Lianne A. Hulshof, Elly M. Hol, Wytse J. Wadman, Davide Iannuzzi

**Affiliations:** Department of Physics and Astronomy and LaserLaB, VU Amsterdam, The Netherlands; Department of Translational Neuroscience, Brain Center Rudolf Magnus, University Medical Center Utrecht, The Netherlands; Netherlands Institute for Neuroscience, An Institute of the Royal Netherlands Academy of Arts and Sciences, The Netherlands; Center for Neuroscience, Swammerdam Institute for Life Sciences, University of Amsterdam, The Netherlands

**Keywords:** viscoelasticity, indentation, structure-stiffness relationship, brain mechanics

## Abstract

There is growing evidence that mechanical factors affect brain functioning. However, brain components responsible for regulating the physiological mechanical environment and causing mechanical alterations during maturation are not completely understood. To determine the relationship between structure and stiffness of the brain tissue, we performed high resolution viscoelastic mapping by dynamic indentation of hippocampus and cerebellum of juvenile brain, and quantified relative area covered by immunohistochemical staining of NeuN (neurons), GFAP (astrocytes), Hoechst (nuclei), MBP (myelin), NN18 (axons) of juvenile and adult mouse brain slices. When compared the mechanical properties of juvenile mouse brain slices with previously obtained data on adult slices, the latter was ~ 20-150% stiffer, which correlates with an increase in the relative area covered by astrocytes. Heterogeneity within the slice, in terms of storage modulus, correlates negatively with the relative area of nuclei and neurons, as well as myelin and axons, while the relative area of astrocytes correlates positively. Several linear regression models are suggested to predict the mechanical properties of the brain tissue based on immunohistochemical stainings.

## 1. Introduction

There is an increasing interest in the mechanical properties of the brain due to the emerging role of physiological mechanical environment in normal brain functioning and involvement of mechanics in disease progression [1]. At a single cell scale, mechanical cues have been showed to regulate brain development [2], stem cell differentiation [1] and morphology of brain cells [3, 4]. For instance, axons of neurons have been shown to grow towards softer tissue *in vivo* [5], and adapt their morphology and stiffness to the rigidity of the substrate in *vitro* [6]. On a tissue scale, there is growing evidence that changes in brain tissue architecture that occurs during neuropathophysiological processes, development, or physiological aging affect the mechanical properties of the brain and thus the local mechanical environment of neurons and glia. To mention a few, reduction of shear modulus was observed during neuroinflammation [7], Alzheimer’s disease [8, 9], multiple sclerosis [10, 11, 12], glial scaring [13] and cancer [14, 15].

Despite the growing evidence of involvement of mechanical cues in brain functioning, there is a lack of fundamental understanding of how different brain components contribute to the overall stiffness of the brain. Anatomical regions of the brain differ in their structural composition, from white-matter (WM) regions composed of fiber bundles with varying degree of myelination and thickness to gray-matter (GM) regions with various densities of neurons, glia, and their arborizations. It is thus not surprising that the different brain regions have been reported to have heterogeneous mechanical properties [16, 17, 18, 19, 20, 21, 22, 23, 24, 25, 26, 27, 28] and that the relationship between some of the components and stiffness was indicated [13, 17, 29]. Despite this body of work, how the mechanical properties and structural composition of the brain relate to each other remains elusive.

As multiple brain components are present within each brain region, measurements of mechanical properties while the composition of the brain changes could indicate which brain components regulate the mechanical environment. One of the naturally occurring modifications of the brain tissue structure is postnatal brain development. During this process, the brain undergoes structural changes such as maturation of extracellular matrix (ECM), myelination, decrease in water content and cell number, dendritic pruning and synaptogenesis, all of which might be accompanied by mechanical alterations [29, 30, 31, 32, 33, 34]. A majority of previous studies have already reported that stiffness increases with maturation [25, 27, 28, 35], yet direct correlations with structural components of measured regions were never investigated. Therefore, the co-quantification of mechanical properties and the composition of the developing brain not only would shed light on structure-stiffness relationship of the brain but also on postnatal maturation of the brain.

In this study, we used a depth-controlled oscillatory indentation technique to map the mechanical properties of individual regions of the hippocampus and cerebellum of horizontal brain slices extracted from juvenile mice. The selected indentation profile enabled viscoelastic characterization in terms of storage and loss moduli, which corresponds to elastic and viscous responses of the material to deformation, respectively, at the tissue scale and physiologically relevant oscillation frequency. Previously, we used the same indentation protocol to map the mechanical properties of adult mouse brain slices, showing that the mechanical properties resemble anatomical regions and that high cell density regions are softer than regions with low cell density, although, the contribution of other brain components were not addressed [36]. In the present study, we quantified relative area A_*rel*_ covered by immunohistochemical staining of NeuN (nuclei of neurons), GFAP (astrocytes), Hoechst (all cell nuclei), MBP (myelin), NN18 (axons) of juvenile and adult mouse brain sections. Differences in terms of storage modulus and stained components between adult and juvenile mouse hippocampus are discussed, as well as correlations between them.

## 2. Methods

### 2.1. Sample preparation for indentation

Two age groups of WT mice (C57Bl6/Harlen) were used for indentation experiments: juvenile (1 month old) and young adult (6 and 9-month-old), (for the latter, we refer the reader to [36]. All experiments were performed in accordance with protocols and guidelines approved by the Institutional Animal Care and Use Committee (UvA-DEC) operating under standards set by EU Directive 2010/63/EU. All efforts were made to minimize the suffering and number of animals. The mice were decapitated, the brain was removed from the skull and stored in ice-cold carbonated artificial cerebrospinal fluid (ACSF). Slices were cut in a horizontal plane with a thickness of approximately 300 μm using a VT1200S vibratome (Leica Biosystems, Nussloch, Germany). Slices from 3 to 4 mm of dorsal-ventral positions of the hippocampus were selected for the measurements, where composition along the thickness can be considered homogeneous. After 1 h rest time, single brain tissue slice was placed in a perfusion chamber coated with 0.1% polyethylenimine for adherence between the glass slide and brain slice, stabilized with a harp and supplied with carbogen saturated ACSF solution. Indentation measurements were performed within 8 h.

### 2.2. Dynamic indentation setup and measurement protocol

Operation of the setup, including ferrule-top force transducer [37, 38] for indentation measurements, has been previously described [36]. Indentation mapping was performed in parallel lines with the distance between two adjacent locations of 50 μm, which assured that deformed areas do not overlap. Measurements were carried out on slices from eight mice for the hippocampus and on six mice for cerebellum with the total number of measurement points *n* = 1704 and *n* = 380, respectively. Previously published oscillatory ramp data on adult mouse brain originated from five mice with the total number of measurement points *n* = 1034 [36].

Ferrule-top probes of 0.2-0.5 N/m stiffness and 60-105 μm bead radius were selected for these experiments to have enough resolution to sample individual regions of the brain but also to be able to indent deeper than the surface roughness of the sliced brain tissue which was previously reported to be 1-3 μm [22]. Indentation-depth controlled oscillatory ramp profile consisted of 0.2 μm amplitude oscillations at 5.62 Hz frequency superimposed on top of a loading ramp at an approximate strain rate of 0.01. The indentation-controlled feedback was triggered at an approximate load of 15 nN, which resulted in the initial uncontrolled 1-3 μm indentation depth, later corrected in post-processing procedures.

### 2.3. Indentation data analysis

The raw data was analyzed with custom-written MATLAB functions [36]. Storage *E*′ and loss *E*’’ moduli were calculated according to [39]:

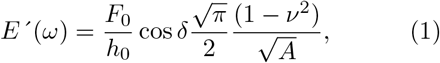

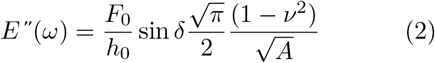

where *ω* is the frequency, *F*_0_ and *h*_0_ are the amplitudes of oscillatory load and indentation-depth, respectively, *δ* is the phase-shift between indentation and load oscillations, *v* is the Poisson’s ratio (0.5 assuming incompressible material), *A* = *πa*^2^ is the contact area. The contact radius *a* was estimated as 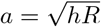 where *h* is the indentation depth and *R* is indenter tip radius.

Indentation depth was converted to the averaged strain according to *ε* = 0.2 × *a/R* [40], thus mechanical properties were measured in the range between 5 and 8% strain, which fulfills small strain approximation[41]. While contact adhesion was observed as a pull-off force during retraction, it was not taken into account, as we assume that, under deep indentation conditions, the nominal area dominates over the actual one. Finally, all indentation curves were checked visually to remove curves where either indentation started in contact or measurements were disturbed by external noise.

### 2.4. Imaging of 300 μm thickness slices

An inverted microscope (Nikon TMD-Diaphot, Nikon Corporation) was used to image the slice during the measurements with a 2 × magnification objective (Nikon Plan 2X, Nikon Corporation). Images were recorded with a CCD camera (WAT-202B, Watec). After the measurements, the slices were fixed in 4% PFA overnight at 4 °C. The slices were stained with Hoechst to label cell nuclei and imaged with Leica DMRE fluorescence microscope (Leica Microsystems, Wetzlar, Germany). Images of the live slices with indentation locations were overlapped over corresponding fluorescent images and each indentation location was assigned to an anatomical region.

### 2.5. Imaging and immunohistochemistry of 30 μm thickness slices

Two age groups of WT mice (C57Bl6/J) were used for immunohistochemistry: 1 and 6-month-old. All animals were housed under standard conditions with ad labium access to water and food. All experiments were approved by the Animal Experimentation Committee of the Royal Netherlands Academy of Arts and Sciences (KNAW) and in accordance with the European Community Council Directive of 24 November 1986 (86/609/EEC).

Mice were anesthetized with 0.1 ml Euthanimal 20% (Alfasan 10020 UDD) and transcardially perfused with 1X PBS. Brains were removed and collected in 4% paraformaldehyde for 48 hours before being transferred to 30% sucrose with sodium azide and stored at 4°C. Before cutting, brains were snap-frozen in isopenthane and mounted using Tissue-Tek (Sakura). Using a cryostat, brains were sliced horizontally in 30 μm thick slices and collected in 1X PBS, which was then replaced by cryopreservation medium (19% glucose, 37.5% ethylene glycol in 0.2 M PB with sodium azide) and stored at −20°C until further processing.

Slices were washed 3 times with PBS before they were blocked with 10% Normal Donkey Serum (NDS, Jackson ImmunoResearch, 017-000-121) and 0.4% Triton-X in 1X PBS for one hour at RT. Sections were incubated with different primary antibodies (see Table S4) diluted in 200 μl 10% NDS and 0.4% Triton-X blocking medium ON at 4°C. Thereafter, they were washed 3 times with 1X PBS and then incubated with 1:1000 secondary antibodies or 1:500 Wisteria floribunda agglutinin (WFA) dye diluted in 200 μl 3.3% NDS and 0.13% Triton-X in 1X PBS ON at 4°C, washed 3 times with 1x PBS and stained with 1:1000 Hoechst dissolved in 500 μl 1x PBS for 10 min at RT. Slices were washed 2 times with 1X PBS and 1 time with MilliQ before mounting on microscope slides using Mowiol (10% Mowiol (Millipore, 475904), 0.1% diazabicyclo(2,2,2)-octane, 0.1 M Tris and 25% glycerol in H_2_O; pH 8.5).

Imaging was done with Zeiss Axioscope.A1 epimicroscope operated with AxioVision software, using 10x Plan-NeoFluar objective.

### 2.6. Image analysis

To compare composition of different anatomical regions, fluorescent images were converted to black and white images by using a threshold (see Fig. S4 for thresholded images). Relative area fraction A_*rel*_ of manually identified regions was calculated for each stained component:

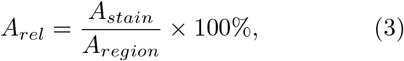

where A_*stain*_ is area covered with stained component and A*_region_* is total area of the region. Images of 6-8 slices from 2-3 animals were used for quantification of each component. The image analysis was performed using Fiji (ImageJ).

### 2.7. Linear regression analysis

Linear models were generated by using the *stepwiselm* MATLAB function. The function starts with the constant model and adds parameters when the predictive power of the model is statistically higher with the parameter than without it (evaluated based on the change in the sum of squared errors; p-values from F-tests).

### 2.8. Statistical analysis

The normality of data distribution was tested with the Shapiro-Wilk test. In the case of a normal distribution, statistical differences between independent groups were investigated with the two-sample t-test while the Wilcoxon rank-sum test was used for non-normally distributed data. All statistical analyses were performed with Statistics and Machine Learning Toolbox (version 2017a, The Mathworks, Natick, MA, USA).

## 3. Results

### 3.1. Mechanical properties of juvenile mouse brain: hippocampus and cerebellum

To characterize local mechanical properties of hippocampus and cerebellum, dynamic indentation mapping was performed on acute mouse brain slices (see Methods 2.1) at 50 μm resolution by indenting with an oscillating ramp at 5.6 Hz frequency up to 8% strain (see Methods 2.2). The viscoelastic properties were quantified in terms of storage *E*’ and loss *E*’’ moduli, and damping factor tan(*δ*), which is the ratio between loss and storage modulus (see Methods 2.3). The brain slices were imaged during dynamic indentation mapping to identify measured anatomical regions (Methods 2.4). Fig. 1 shows two examples of maps of storage modulus of the hippocampus, subregions of DG, CA3 and CA1 (Fig. 1 A, B) and two examples of maps of cerebellum (Fig. 1 C, D) where lighter color indicates stiffer tissue and darker color indicates softer tissue. Contrast due to mechanical heterogeneity agreed with the shapes of anatomically defined brain regions.

**Figure 1:**
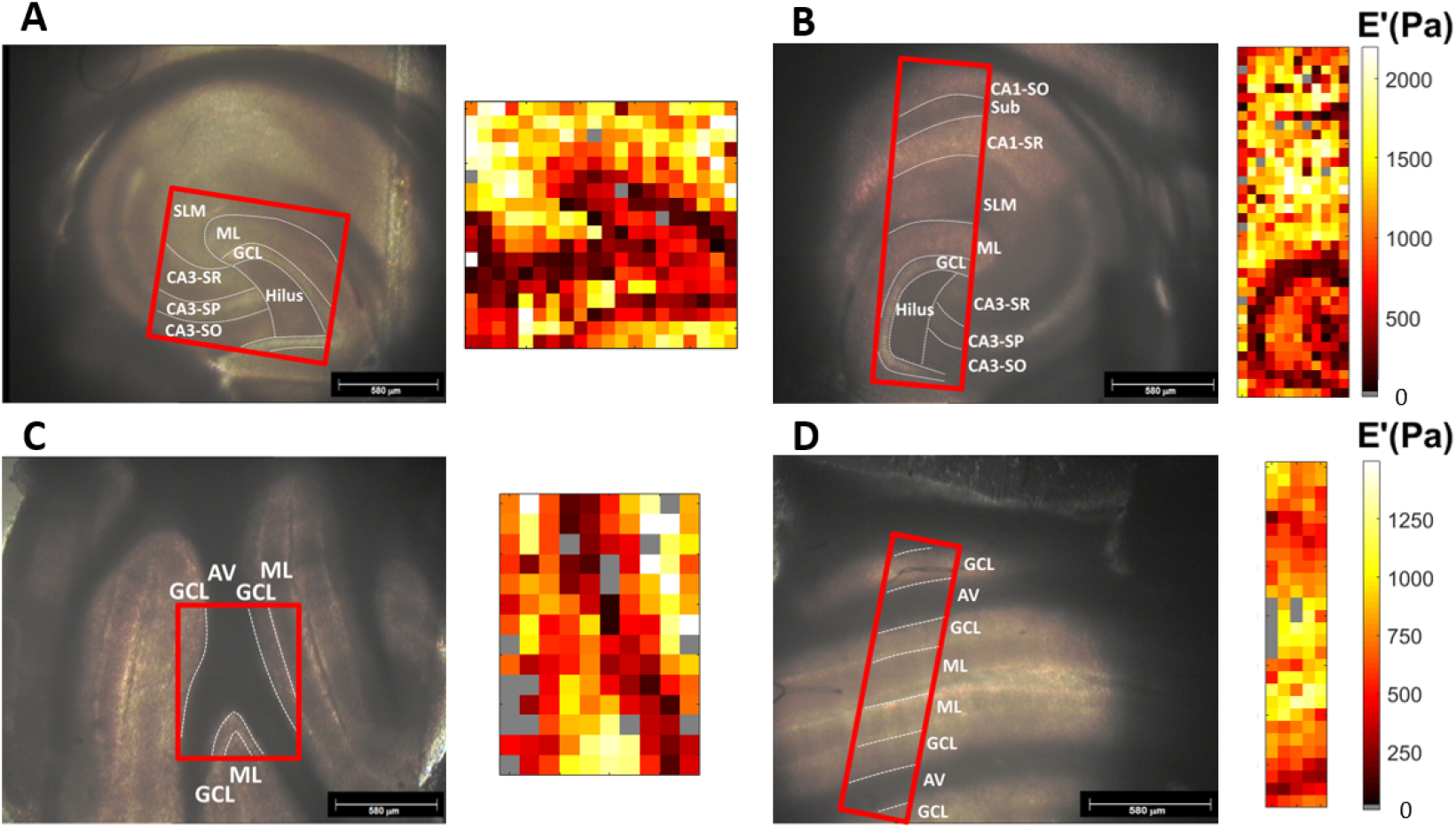
Microscope images of (A, B) hippocampus and (C, D) cerebellum where area measure by indentation is indicated with red rectangular. Next to it, maps of storage modulus *E*’ at a 50 μm × 50 μm resolution obtained with 0.2 μm oscillation amplitude, 5.62 Hz oscillation frequency, and at 7% strain. Dashed lines indicate boundaries of anatomical regions. Abbreviations: Alv - alveus, Sub - subiculum, SLM - stratum lacunosum moleculare, SR - stratum radiatum, SP - stratum pyramidale, SO - stratum oriens, ML - molecular layer, GCL - granule cell layer, AV - arbor vitae; large regions: DG - dentate gyrus, CA - cornus ammonis. Gray color indicates failed measurements.

To account for the inter-animal variation of mechanical measurements, the same indentation protocol was repeated on slices from multiple animals. Data from different slices were merged for each anatomical region (Fig. 2 A, B). The storage modulus of the hippocampus and cerebellum were mechanically heterogeneous with averaged storage modulus values *E*’ at 7% strain varying from 0.4 to 0.9 kPa in the cerebellum and from 0.4 to 1.6 kPa in the hippocampus (Fig. 2 (A)). The averaged damping factor tan(*δ*) was higher in the cerebellum (0.56-0.69) than in the hippocampus (0.42-0.55) (Fig. 2 B), indicating higher energy dissipation potential of cerebellum. Fig. 2 (C) shows a color-coded reconstruction of the storage modulus *E*’ of hippocampus and cerebellum, based on the averaged results in Fig. 2 (A), and the fluorescence images of the same brain areas stained for nuclei and myelin with identified anatomical regions.

**Figure 2:**
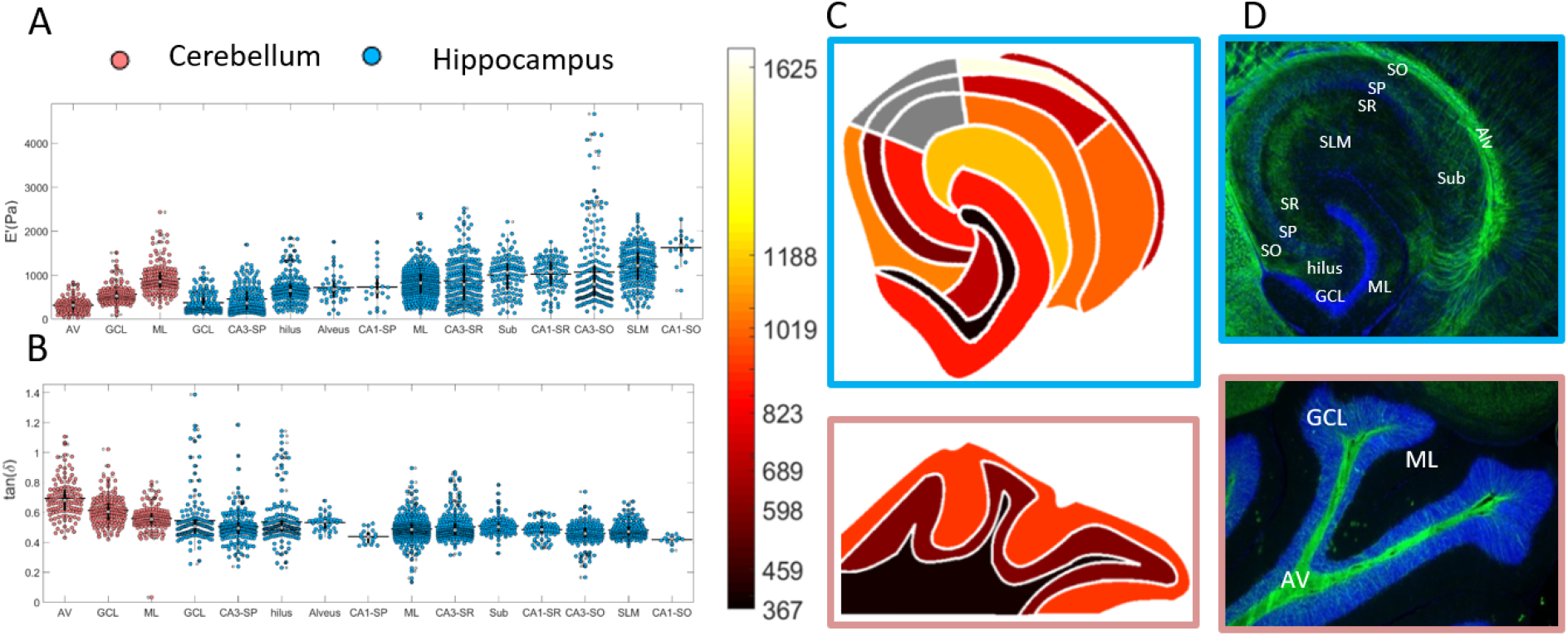
A) Storage modulus *E*’ and B) damping factor tan(*δ*) of regions in cerebellum (red) and hippocampus (blue) ordered from softest to stiffest, obtained at 7% strain and 5.62 Hz frequency of oscillations. Horizontal bar is mean value, vertical bar is boxplot with median value marked as white dot. Data is merged from multiple animals (N = 2–8 animals, n = 16–335 indentation measurements per region). C) Visual reconstruction of averaged storage modulus values and D) fluorescent images of hippocampus and cerebellum stained for nuclei (Hoechst) in blue and myelin (MBP) in green.

The oscillatory ramp indentation profile allowed us to investigate the strain-dependent mechanical properties of brain tissue i.e. nonlinearity. All measured brain regions showed an increase in storage modulus *E*’ with strain (see Fig. 3) where stiffening was less pronounced in softer regions (39-191 kPa vs. 234-493 kPa per 1% of the strain, respectively). Furthermore, the spread in averaged values was larger at higher strains (0.2-0.6 kPa at 5% strain and 0.4-1.7 kPa at 8% strain) which means that the contrast in mechanical properties between brain regions is more pronounced at higher strains than at lower strains. The damping factor tan(*δ*) decreased with strain for regions in the hippocampus (0.03-0.08 per 1% strain), while it was rather constant for cerebellum (decrease of 0.01 per 1% strain), which suggests differential viscoelastic behavior of these two regions.

**Figure 3:**
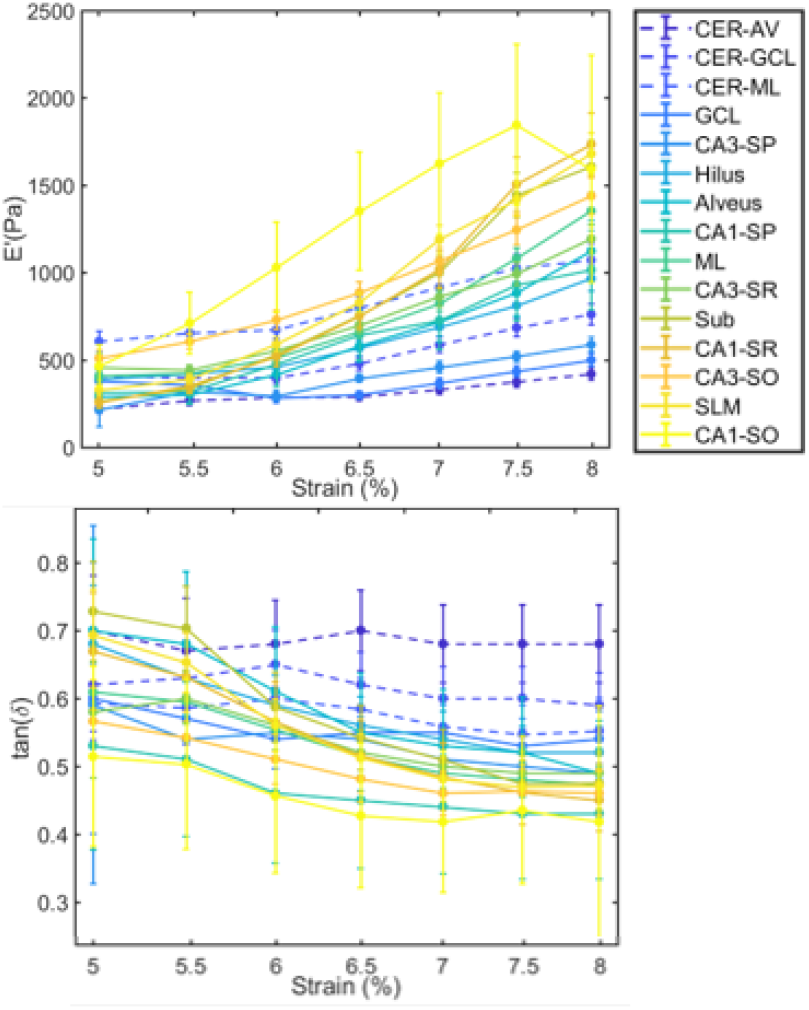
Averaged storage modulus *E*’ and damping factor tan(*δ*) as a function of strain for different brain regions, identified in the legend. Dashed lines for cerebellum regions and solid lines for hippocampal regions.

### 3.2. Adult brain is stiffer than juvenile brain

To compare mechanical properties of juvenile and adult mouse brain, the mechanical data from previously obtained hippocampal regions of the adult mouse brain [36] was plotted together with the matching regions of the juvenile brain (see Fig. 4A), where sample preparation and measurement protocols were identical. The averaged storage moduli *E*’ of adult mouse brain were significantly higher than the storage moduli of juvenile mouse brain for all measured regions, with the only exception of CA3-SP region, where the difference was not significant. The increase in storage modulus between juvenile and adult mouse brain was low for GCL, CA3-SP and Alveus (20-50%) which are relatively soft regions and densely packed with either neurons or fibers. Stiffer and less packed regions such as ML and SLM had a higher increase in stiffness with age (60-150%, see above Fig. 4). The damping factor tan(*δ*) was similar when comparing adult and juvenile mouse brains (juvenile ~0.49±0.05 and adult ~o0.50±0.0I, see Table S2). These results suggest that during maturation of mouse brain tissue, stiffness of the hippocampus substantially increases while tan(*δ*) remains constant.

**Figure 4:**
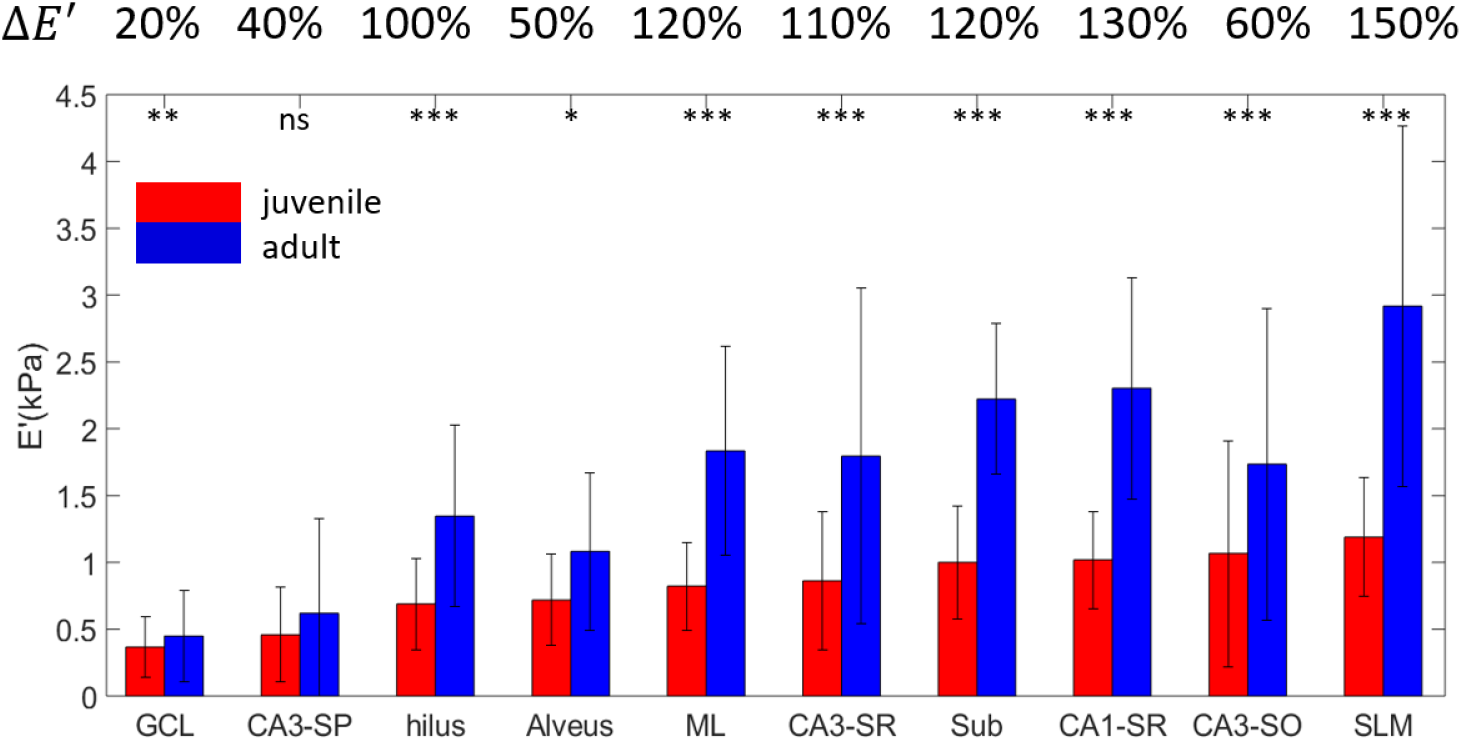
Comparison of averaged storage modulus *E*’ of juvenile (red) and adult (blue) hippocampal subregions. Percentages above the graph are relative increase in storage modulus with age. Error bars are standard deviation. ***p<0.001, **p<0.01, *p<0.5, ns - non significant.

### 3.3. Structure-stiffness relationship between brain regions

To assess structural differences between measured brain regions, and between juvenile and adult mouse brains, we performed immunohistochemical stainings to label main brain components: all cell nuclei (Hoechst), nuclei of neurons (NeuN), astrocytes (GFAP), axons (NN18), myelin (MBP), perineuronal nets (WFA) and dendrites (MAP2) (see Fig. 5 A, staining procedure in Methods 2.5). The amount of each component was quantified as A_*rel*_(%), a relative area covered by the stained component within each region (Fig. 5 B, see Table S3 for all A_*rel*_ values and Methods 2.6 for protocol). MAP2 and WFA stainings were compared qualitatively because they showed less conspicuous regional differences (Fig. S3).

**Figure 5:**
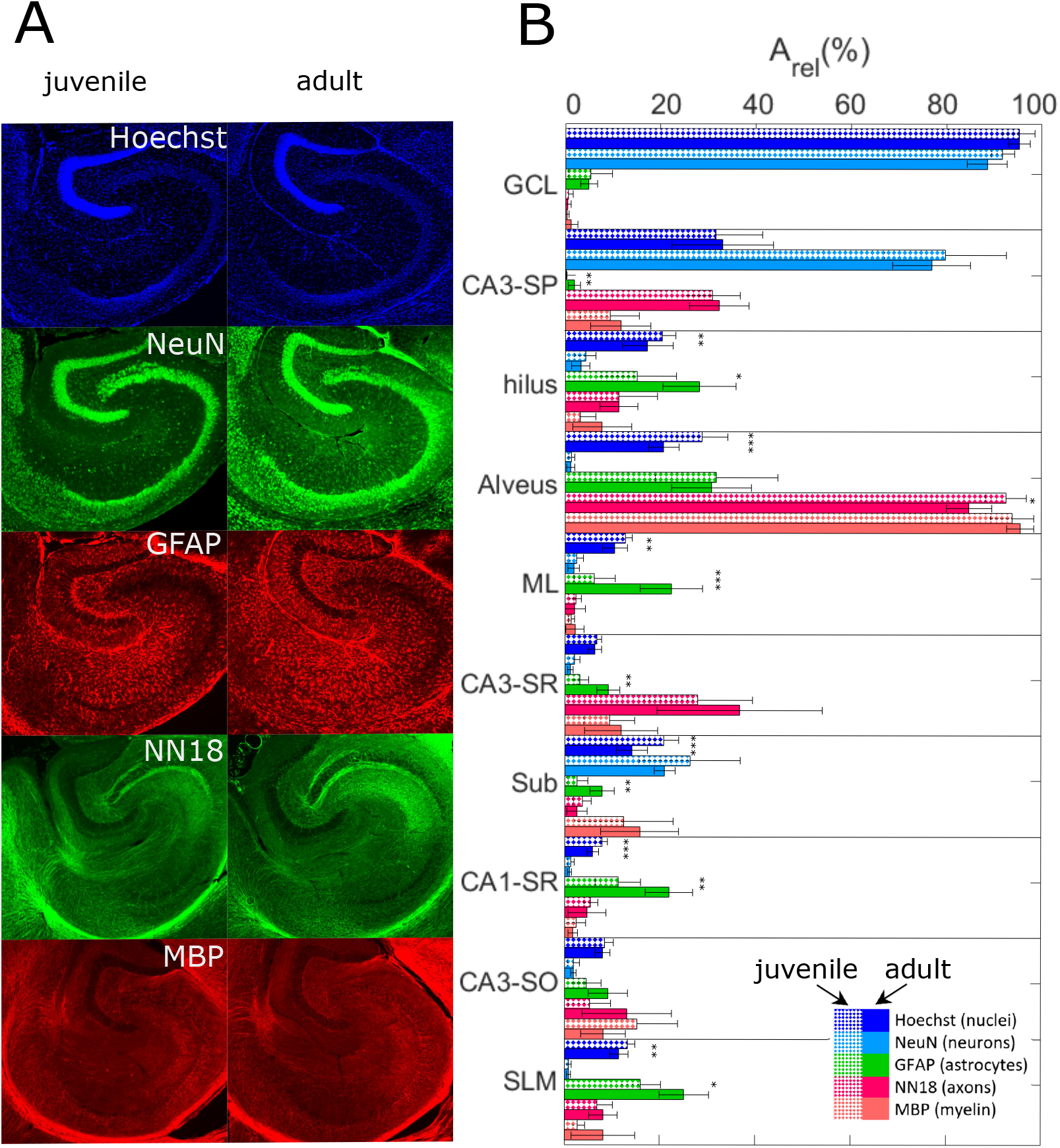
Comparison of immunofluorescent images of different brain components between juvenile and adult mice hippocampus. A) Representative fluorescent images of immunohistochemical stainings of all cell nuclei (Hoechst), nuclei of neurons (NeuN), astrocytes (GFAP), axons (NN18) and myelin (MBP). Dimensions of images are 2065 μm (width) and 1878 μm (height). B) Relative area A_*rel*_ (%), averaged over multiple slices obtained from 2-3 animals per age group, of different brain components of juvenile (first bar) and adult (second bar) hippocampal mouse brain regions. Statistical differences in terms of p-values (see Table 4). ***p<0.001,**p<0.01, *p<0.5.

When comparing juvenile and adult mouse brains, a significant increase in A_*rel*_ of GFAP was observed for most of the regions (Fig. 5 B, Table S4). Moreover, A_*rel*_ of nuclei significantly decreased in many regions. Although A_*rel*_ of other quantified components, i.e. NeuN, NN18 and MBP, showed either decrease or increase with age, differences were not significant (see Fig. 5 B). Interestingly, increase in storage modulus from juvenile to adult brain (Δ*E*’) had moderate positive correlation with the increase in A_*rel*_ of GFAP (Pearson’s correlation coefficient r = 0.61, p = 0.06, Fig. 6 F) while correlations were weak for all other components (r = −0.18 for Hoechst; 0.4 for NeuN; 0.05 for NN18; 0.25 for MBP).

**Figure 6:**
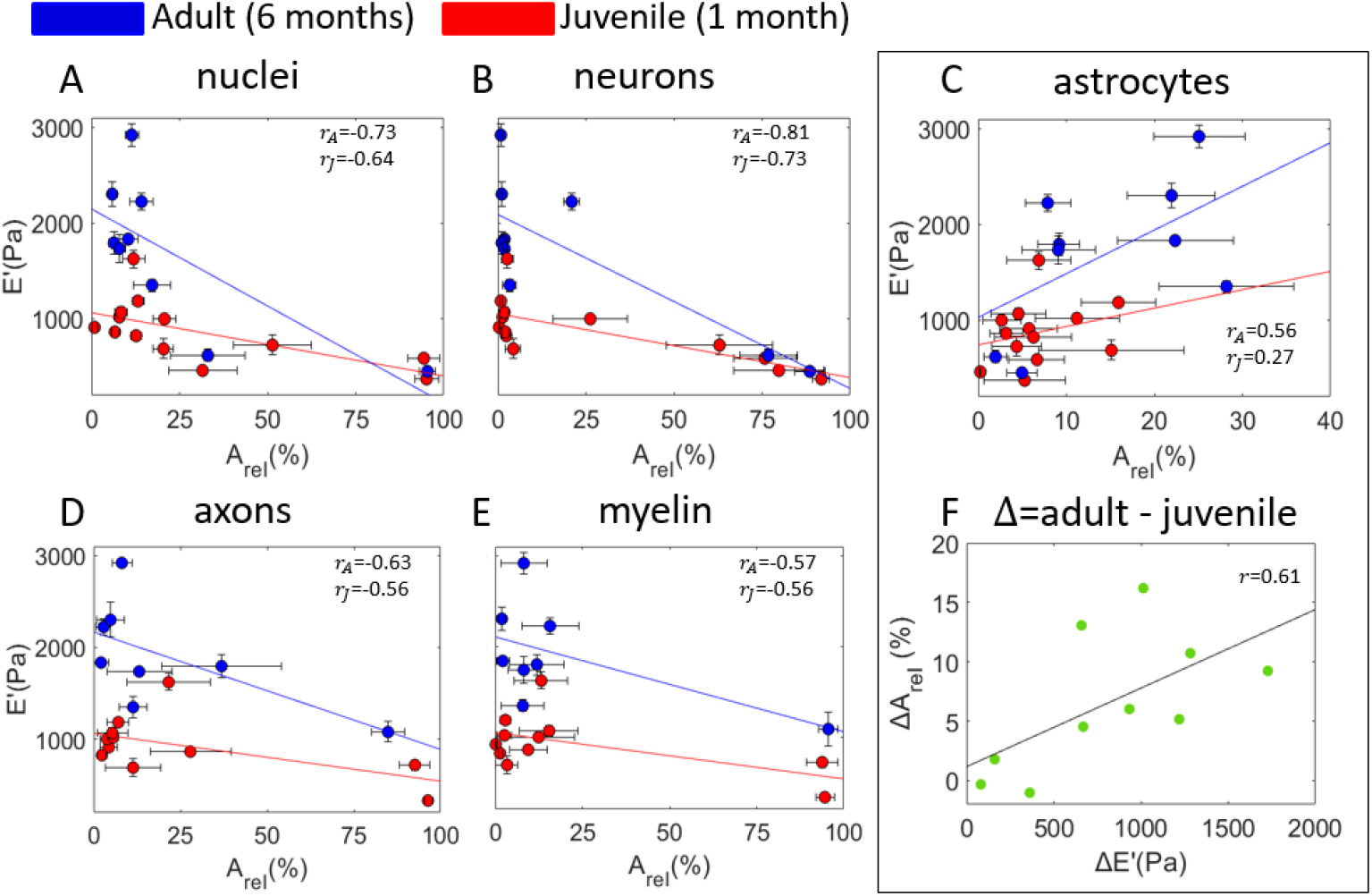
Storage modulus *E*’ as a function of relative area A_*rel*_ covered by A) all nuclei, B) nuclei of neurons, C) astrocytes, D) axons and E) myelin of measured regions, both juvenile (red) and adult (blue). WM regions are excluded in A-C) plots while high-density GM regions are excluded in C-D) plots. Pearson’s correlation coefficient identified above, r_*a*_ for adult data and r_*j*_ for juvenile. F) Increase in A_*rel*_ of astrocyte staining as a function of the increase in storage modulus *E*’ when comparing juvenile and adult data. Pearson’s correlation factor r=0.61 (p=0.06).

A vast number of WFA positive cells were in Subiculum while a few cells were WFA positive in the SP layer of CA1-CA3 regions. Comparison between juvenile and adult mouse brains showed a clear increase in perineuronal nets (PNNs) positive cells in Subiculum with age (Fig. S3). Staining with MAP2 of dendrites showed organizational rather than quantitative differences between regions (Fig. S3). Dendrites appeared as a honeycomb structure around cells, parallel and long in CA1-SR and Sub regions and homogeneous in all of the other regions. A comparison between adult and juvenile did not show observable differences in MAP2 staining.

Assessment of mechanical and structural regional differences was done by plotting storage modulus *E*’ as a function of A_*rel*_ of different brain components, both adult (in blue) and juvenile (in red) (see Fig. 6). As both, high-density cell regions and high-density fiber bundles (WM), are mechanically soft, correlation analysis for nuclei, neurons and astrocytes (Fig. 6 A-C) was done by excluding WM regions (Alv and AV) while correlation analysis for myelin and axons (Fig. 6 D, E) was done by excluding regions with high-density of nuclei (GCL, SP3 and SP1, A_*rel*_ > 50%). As a result, *E*’ was found to correlate negatively with A*_*rel*_* of nuclei (Pearson’s correlation factor for adult r_a_=-0.73 and for juvenile r_*j*_=-0.64), neurons (r_*a*_=-0.81, r_*j*_=-0.73), axons (r_*a*_=-0.57, r_*j*_=-0.56) and myelin (r_*a*_=-0.63, r_*j*_ =-0.56). Moreover, storage modulus *E*’ correlated positively with A*_*rel*_* of astrocytes (r_*a*_=0.56, r_*j*_ =0.27). Together, these results show that relatively cell-free and axon-free regions are the stiffest while regions that are tightly packed with either nuclei or axons are the softest, whereas a higher number of astrocytes seemed to be responsible for higher stiffness values between regions and when comparing juvenile and adult brains.

### 3.4. Linear regression model for storage modulus prediction from immunohistological data

Previous studies have shown that the mechanical properties of spine tissue can be predicted from histological data [42, 23]. We applied linear regression analysis (see Methods 2.7) to investigate which of the measured histological parameters are needed for the prediction of storage modulus values of individual brain regions. The best prediction (R^2^=0.60) of storage modulus E’ of juvenile brain areas, when including indentation data of hippocampus and cerebellum, was achieved with the relative area of NeuN A*_NeuN_* (p=0.002) and GFAP A_*GFAP*_ (p=0.01) stainings:

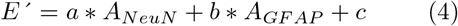

where *a* = −7.5 ± 1.9, *b* = −14.9 ± 5.0 and *c* = 1172 ± 103 are linear regression model parameters.

The best prediction (R^2^=0.70) of storage modulus E’ of adult brain hippocampal regions was achieved with the relative area of NeuN A*_NeuN_* (p=0.007) and NN18 A_*NN*18_ (p=0.07) stainings:

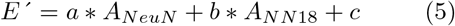

where *a* = −17.9 ± 4.8, *b* = −13.0 ± 6.2 and *c* = 2230 ± 220 are model parameters.

To test whether including other histological parameters would improve the prediction of regional mechanical properties, estimations of densities of cells, neurons, glia, oligodendrocytes, astrocytes and microglia of adult hippocampal regions were acquired from Blue Brain Cell Atlas (BBCA) (for more information [43]). It is important to note that these estimations of cell densities are calculated from the entire volume of the brain region whereas we measured mechanical properties only at the specific plane within the brain. Therefore, by using this data we assumed that there are no variations in cell densities and mechanical properties within the volume of the brain areas. As a result, linear regression model revealed that density of all cells P_*allcells*_ (p=5.4×10^−6^), density of glia P_*glia*_ (p=0.01) and relative area of NN18 A_*NN*18_ (p=0.002) could give the prediction of storage modulus values of hippocampal areas with the highest R-squared value (R^2^=0.96):

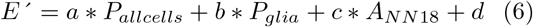

where *a* = −0.004 ± 0.0004, *b* = −0.0002 ± 0.0006, *c* = −13.8 ± 2.5 and *d* = 2489 ± 140 are model parameters.

## 4. Discussion

### 4.1. Viscoelastic mapping of hippocampus and cerebellum

In this study, we selected 50 μm mapping resolution and indentation depth of up to 10-17 μm (8% strain) to obtain mechanical properties of individual regions of hippocampus and cerebellum of the juvenile mouse brain tissue (Fig. 1). We found that both the hippocampus and the cerebellum were mechanically heterogeneous where mechanically distinct regions matched morphologically defined anatomical regions. Similar relative mechanical differences between hippocampal subregions of juvenile mouse brain tissue were found in the previous study on adult mouse brain [36] where the measurement protocol was identical to the one in this study. Other studies have been done at different measurement scales and indentation protocols, making it difficult to compare results between studies. For example, some of the studies used tips with the radius of 250 μm resulting in contact area much larger than individual anatomical regions of hippocampus and thus measuring averaged mechanical properties over multiple layers [16, 17, 18, 19, 25, 26, 27, 22, 44]. Other studies used smaller tips and indentation depths (R < 20 μm, h < 4ιm [28, 45, 46]), where reported relative differences between regions do not agree with our findings.

In comparison to studies regarding mechanical properties of the cerebellum, we were able to differentiate between three layers (Fig.2 A), two stiffer GM regions (ML and GCL) and softer WM region (AV), while previous studies only differentiated between GM and WM with contradicting results between the studies. Previous quasi-static indentation measurements with small spherical tips (R = 10-20 μm) agreed with our findings in cerebellum, where GM was stiffer than WM [22, 47], while stress relaxation measurements, performed with flat cylindrical punch of 250]im radius, have found that GM was softer than WM [26] or showed no significant differences [25, 27, 48, 49].

It is well-known that the mechanical behavior of the brain tissue is nonlinear and viscoelastic and thus we selected an oscillatory ramp indentation protocol which allowed us to obtain multiple viscoelastic parameters of the brain, i.e. strain-dependent storage modulus *E*’ and damping factor tan(*δ*) (see Fig. 3). These measurements revealed that the cerebellum and the hippocampus have very different viscoelastic properties, i.e. hippocampus showed higher stiffening with strain and had lower damping ratio at higher strains when comparing with the cerebellum. We hypothesize that these differences in mechanical behavior could be related to the difference in their location within the brain. While the hippocampus is positioned in the middle of the brain, the cerebellum is located at the back of the brain, close to the skull and thus more vulnerable to the head injury which could also explain its better energy damping capability. These findings demonstrate the potential of measuring multiple mechanical parameters, i.e. storage modulus, damping factor and their depth dependency, as it gives more information about the structure and function of the brain than typical static measurements. For example, a recent study has shown that stiffening with increasing compression is a hallmark for the differentiation between healthy and glioma brain tissue [50].

Overall, this data can be used to support future biomechanical and biochemical studies. To mention a few, local native mechanical properties of brain tissue are needed when culturing neurons and glial cells on compliant substrates [3, 4, 51, 52, 53], when designing mechanically compatible brain implants [54] and modeling traumatic brain injuries [55].

### 4.2. Changes in mechanical properties of the hippocampus with age

Mechanical cues are involved in the regulation of brain development as shown by previous studies[1]. We observed 20-150% increase in stiffness when comparing juvenile and adult mouse hippocampal regions (Fig. 4 A), where densely packed regions, either with cells or fibers, stiffened less than loosely packed regions. Most of the previous studies on rodent brain slices are in agreement with our findings. For example, when compared 17-18 postnatal days (PND) and fully mature rat brains, the latter were found to be stiffer for most of the regions [25, 27]. Another study also found that the elastic modulus of rat hippocampus and cortex increased more than 2 times when comparing prenatal and adult brains [28]. In contrast, one study compared the elastic modulus of different brain regions between 10-20 weeks and 100-105 weeks old mice, and only WM in stratum was significantly stiffer (1.5 fold) in older animals [47]. Measurements on intact brains rather than brain slices have also opposing results. For example, 12 weeks old mice were 13-59% stiffer than 6 weeks old mice [35] while immature rat brains (PND13 and PND17) were stiffer than mature ones (PND43 and PND90) [56]. Therefore, our study contributes to the body of evidence that brain tissue stiffens with maturation.

### 4.3. Changes in immunohistochemically stained components of the hippocampus with age

Co-registration of both, mechanical properties and brain components at different developmental stages, could indicate which structural components are responsible for mechanical maturation of brain tissue. Although many structural changes that take place during the maturation of the brain are known, in many cases the data is obtained from large brain regions rather than individual cell layers and limited to the specific development stage, making it difficult to objectively compare the stiffness and composition of brain tissue between different studies. Here, we performed the direct comparison, i.e. immunohistochemical stainings of different brain components (Fig. 4 D) of adult and juvenile mouse brain slices, obtained from the same brain area as slices used for mechanical measurements.

To asses the differences in the cellular composition of hippocampal regions, we stained all cell nuclei (Hoechst), nuclei of neurons (NeuN) and astrocytes (GFAP). Quantification of the relative area covered by stained component of juvenile and adult mouse hippocampus revealed that A_*rel*_ of GFAP staining significantly increased (1.8-16.2%) with age for most of the subregions, which agrees with the previous findings that GFAP is upregulated with maturation of astrocytes and aging of mice [57, 58, 59, 60]. Furthermore, we observed a significant decrease in A_*rel*_ (0.4-8%) of nuclei staining (Hoechst) and a non-significant decrease in A_*rel*_ (0.3-5%) of neuronal nuclei staining (NeuN) with age for the majority of measured hippocampal subregions. In agreement with our finding, the previous study on mice maturation reported that neuronal and non-neuronal cell nuclei densities decreased in the hippocampus when comparing similar age groups to our study (1 and 4 months) [61].

To evaluate the composition of hippocampal regions in terms of cellular arborizations, we stained axons (NN18), myelin (MBP) and dendrites (MAP2). When comparing myelin staining (MBP) of juvenile and adult mouse brain slices, A_*rel*_ was increased (0.8-5.5%) for most of the hippocampal subregions, although the difference was not significant. In mice, myelination between 1 and 6 months has been reported to take place in the corpus callosum, fimbria and cortex [30, 31, 32, 62]. Although there is no data available of myelination in the mouse hippocampus, it has been reported that, in rats, numbers and distribution of myelinated fibers are the same on day 25 and adulthood [63], which agrees with our finding. Furthermore, A_*rel*_ of axonal staining (NN18) was similar between two age groups for most of the hippocampal regions with the exception of Alveus where it was significantly decreased, and CA3-SR and CA3-SO regions, where A_*rel*_ was substantially increased (14.7% and 15.1%) although the differences were not significant. Dendritic staining (MAP2) showed no qualitative differences between juvenile and adult hippocampus. Because neuronal network outgrow, elongation and branching has been reported to take place in early postnatal stages (P<30), it seems plausible that there are no large structural changes of these networks into the adulthood [64, 65, 66, 67, 68, 69].

Brain tissue also contains ECM which forms a fine macromolecular mesh around cell somata, initial segments of axons, and synapses and consists of collagen type IV, HA, fibronectin, laminin, and proteoglycans [70, 71]. Because ECM in the brain lacks filamentous proteins such as fibrous collagen type I [70, 72], it is not expected that ECM regulates brain tissue stiffness [73, 74]. In support of this hypothesis, recently it has been shown that overexpression of ECM components laminin and collagen IV in glial scars correlates with brain tissue softening [13]. Nevertheless, during the development of the brain, ECM transitions from a juvenile-type matrix to a mature one [33, 34, 75]. To check whether there are differences between 1 and 6 months old hippocampus, we stained perineuronal nets, which are particularly enriched with ECM molecules [76]. The noticeable increase in the number of cells wrapped by PNNs was present only in the Subiculum region and thus does not explain the stiffening of all the regions. Previous studies have shown that besides PNNs, hyaluronan/proteoglycan link protein 1 (HAPLN1) increased in the hippocampus between 1.5 and 6 months age [77] while protein levels of neurocan, brevican and tenascin-R were similar. Another study has shown that levels of aggrecan, versican (GAG*α*), brevican increased between 1 and 6 months while levels of neurocan and versican (GAG*β*) decreased, although the study was done on the whole volume of rat brains [75].

So far, there is very little information regarding the structure-stiffness relationship of the brain tissue during maturation reported in the literature. Only one previous study has shown that an increase in stiffness of the hippocampus with age (P10, P17 and adult) agrees with the decrease in water content and increase in protein and lipid (myelin) content [28]. Another study, although only on white matter, has reported that stiffness tripled while myelin content increased from 58 to 74% when comparing prenatal and postnatal bovine brains [29]. In this study, differences in A_*rel*_ of stained components between juvenile and adult were correlated with differences in storage modulus (Fig. 6 F), where only A_*rel*_ of GFAP staining showed moderate positive correlation, which suggests that GFAP positive cells contribute to mechanical stiffening of hippocampal regions during brain maturation.

One of the limitations of this study is the quantification of the relative area covered by the stained component, which only gives a rough estimate of the brain composition. A more thorough structural analysis could include 3D analysis of brain slices and quantification of size and density of cells, orientation and thickness of cellular arborizations and staining of subtypes of cells. Furthermore, we only investigated structural changes at the tissue scale while changes at cellular/axonal scale might also influence tissue mechanics. Finally, volumetric changes of brain components such as extracellular space or volume fraction of different components could also be important factors governing brain tissue mechanics.

There are two other hypotheses proposed in the literature that explains the cause of brain tissue stiffening with maturation. The first hypothesis implies that, with age, the amount of negatively charged glycosaminoglycans (GAGs) increases [28] resulting in elevated Donan osmotic pressure [78] and thus stiffness, which is similar to the behavior observed in articular cartilage [79, 80]. Another hypothesis is that axons in the brain are under tension [73], and axonal tension might increase during brain tissue transition into adulthood. Whether axonal tension or negative charges drive brain tissue stiffening during maturation should be a topic of further investigation.

### 4.4. Correlation between mechanical properties and immunohistochemical stainings

To explain differential mechanical properties of brain regions within the juvenile and adult mouse brains, we performed correlation analysis between relative area covered by immunohistochemical staining A_*rel*_ and storage modulus *E*’. We found that storage modulus and A_*rel*_ of all nuclei, nuclei of neurons, axons and myelin of different brain regions have moderate to high negative correlation (Fig. 6 A, B, D, E) while A_*rel*_ of astrocytes has low to moderate positive correlation, with stronger correlation factor for adult than juvenile in all cases. Based on these results, we hypothesize that the loss of myelin as in demyelinating diseases or loss of neurons as during brain development and aging, and increase in the number of astrocytes as in neuroinflammatory diseases would all result in stiffening of the brain region.

Only one previous study correlated myelin content with the stiffness by comparing different cerebral white matter regions in the bovine brain (myelin content 64-89%), including pre-natal and post-natal (55-89%) brain [29]. However, this study did not include gray matter regions or other brain components. Regarding other CNS tissues, it has been shown that cell density and stiffness correlates positively in retinal and spinal cord tissues where contradiction in comparison to our study might be due to differences in CNS tissue composition or much smaller measurement scale (indentation depth <3.5 μm) [81, 23].

### 4.5. Linear model for predicting mechanical properties of the brain

Prediction of mechanical parameters from histological stainings would allow assessing information about brain stiffness without the need for experimental testing. From the linear regression analysis, we were able to identify that storage modulus could be predicted by using the relative area of NeuN and GFAP staining for the juvenile brain and NeuN and NN18 staining for the adult brain. However, the R-squared values of prediction were only 0.6 and 0.7, respectively, questioning the reliability of such a linear model. The prediction of storage modulus of adult mouse hippocampal regions was improved to R-squared of 0.96 by including the density of all cells and glia from Blue Brain Cell Atlas together with the relative area covered by axons. Regarding data from BBCA, the density of all cells was obtained by Nissl staining for all cell bodies. Furthermore, glial staining from BBCA included oligodendrocytes (CNP and MBP), astrocytes (S100b, GFAP and ALDH1L1) and microglia (TMEM119). Therefore, including other parameters describing tissue composition could improve the predictive power of the model. Future studies should investigate which histological parameters are most relevant for describing the structure-stiffness relationship of the brain tissue by expanding it to other brain regions.

## 5. Conclusions

Dynamic indentation mapping of hippocampus and cerebellum of juvenile mouse brain revealed that viscoelastic parameters vastly differ between individual brain layers. Furthermore, juvenile brain was found to be significantly softer than adult brain. Finally, structure-stiffness relationship of the brain tissue was shown be correlating mechanical differences with the amount of brain components.

## Acknowledgements

The research leading to these results has received funding from the European Research Council under the European Union’s Seventh Framework Programme (FP/2007-2013)/ERC grant agreement no. [615170] and Foundation for Fundamental Research on Matter (FOM), which is financially supported by the Netherlands Organisation for Scientific Research (NWO). The authors further thank M.Marrese and S.V.Beekmans for fruitful discussions.

## Supplementary Material

**Table 1:** Antibodies used for immunohistochemical staining.

| Primary antibody | Dilution | Source (article nr) | Type | Secondary Antibody | Dilution | Source |
| --- | --- | --- | --- | --- | --- | --- |
| Fluorescent immunohistochemistry |  |  |  |  |  |  |
| Hoechst | 1:1000 | Invitrogen, 11534886 | NA | NA | NA | NA |
| Mouse-anti-NeuN | 1:500 | Sigma-Aldrich MAB377 | monoclonal | Donkey-anti-Mouse Alexa Fluor 594 | 1:1000 | Jackson ImmunoResearch 715-585-150 |
|  |  |  |  | Donkey-anti-Mouse Alexa Fluor 488 |  | Jackson ImmunoResearch 715-546-150 |
| Rabbit-anti-GFAP | 1:1000 | CiteAb Z0334 | polyclonal | Donkey-anti-Rabbit Alexa Fluor 488 | 1:1000 | Jackson ImmunoResearch 711-545-152 |
|  |  |  |  | Donkey-anti-Rabbit Alexa Fluor 594 |  | 594, Jackson ImmunoResearch 711-496-152 |
| Mouse-anti-NN18 | 1:1000 | Sigma N5264 | monoclonal | Donkey-anti-Mouse Alexa Fluor 594 | 1:1000 | Jackson ImmunoResearch 715-585-150 |
| Rat-anti-MBP | 1:1000 | Sigma-Aldrich MAB386 | monoclonal | Donkey-anti-Rat Cy3 | 1:1000 | Jackson ImmunoResearch 712-165-153 |
| Mouse-anti-MAP2 | 1:1000 | BioLegend SMI-52P | monoclonal | Donkey-anti-Mouse Alexa Fluor 594 | 1:1000 | Jackson ImmunoResearch 715-585-150 |
| Fluorescein labeled WFA | 1:500 | Vector Laboratories FL-1351 | NA | NA | NA | NA |
| NA – not applicable |  |  |  |  |  |  |

**Table 2:** Storage modulus and damping factor of measured brain regions of 1 and 6-month old mice.

|  | 1 month old |  |  |  |  |  | 6 months old |  |  |  |  |  |
| --- | --- | --- | --- | --- | --- | --- | --- | --- | --- | --- | --- | --- |
| | E'(Pa) | SEM | tan($\delta$) | SEM | n | N | E'(Pa) | SEM | tan($\delta$) | SEM | n | N |
| ML | 823 | 18 | 0.49 | 0.04 | 335 | 6 | 1835 | 48 | 0.46 | 0.03 | 263 | 5 |
| GCL | 367 | 19 | 0.54 | 0.03 | 146 | 6 | 449 | 31 | 0.44 | 0.04 | 123 | 5 |
| Hilus | 689 | 26 | 0.54 | 0.03 | 173 | 6 | 1347 | 62 | 0.46 | 0.03 | 120 | 5 |
| CA3-SO | 1067 | 61 | 0.46 | 0.03 | 192 | 6 | 1736 | 147 | 0.46 | 0.03 | 63 | 4 |
| CA3-SP | 459 | 23 | 0.50 | 0.05 | 232 | 6 | 620 | 66 | 0.54 | 0.19 | 114 | 4 |
| CA3-SR | 862 | 34 | 0.50 | 0.03 | 232 | 6 | 1795 | 117 | 0.46 | 0.03 | 115 | 4 |
| SLM | 1189 | 31 | 0.48 | 0.02 | 204 | 5 | 2918 | 115 | 0.49 | 0.03 | 138 | 5 |
| CA1-SO | 1624 | 93 | 0.42 | 0.02 | 16 | 2 |  |  |  |  |  |  |
| CA1-SP | 727 | 105 | 0.44 | 0.02 | 19 | 2 |  |  |  |  |  |  |
| CA1-SR | 1019 | 38 | 0.49 | 0.02 | 92 | 4 | 2303 | 125 | 0.58 | 0.05 | 44 | 4 |
| Sub | 1000 | 39 | 0.51 | 0.04 | 116 | 2 | 2222 | 89 | 0.50 | 0.02 | 40 | 3 |
| Alv | 718 | 54 | 0.53 | 0.03 | 40 | 3 | 1082 | 187 | 0.47 | 0.04 | 10 | 2 |
| Cer-ML | 915 | 27 | 0.56 | 0.03 | 149 | 6 |  |  |  |  |  |  |
| Cer-GCL | 587 | 17 | 0.60 | 0.03 | 146 | 6 |  |  |  |  |  |  |
| Cer-Av | 332 | 13 | 0.66 | 0.04 | 85 | 6 |  |  |  |  |  |  |

**Figure 1:**
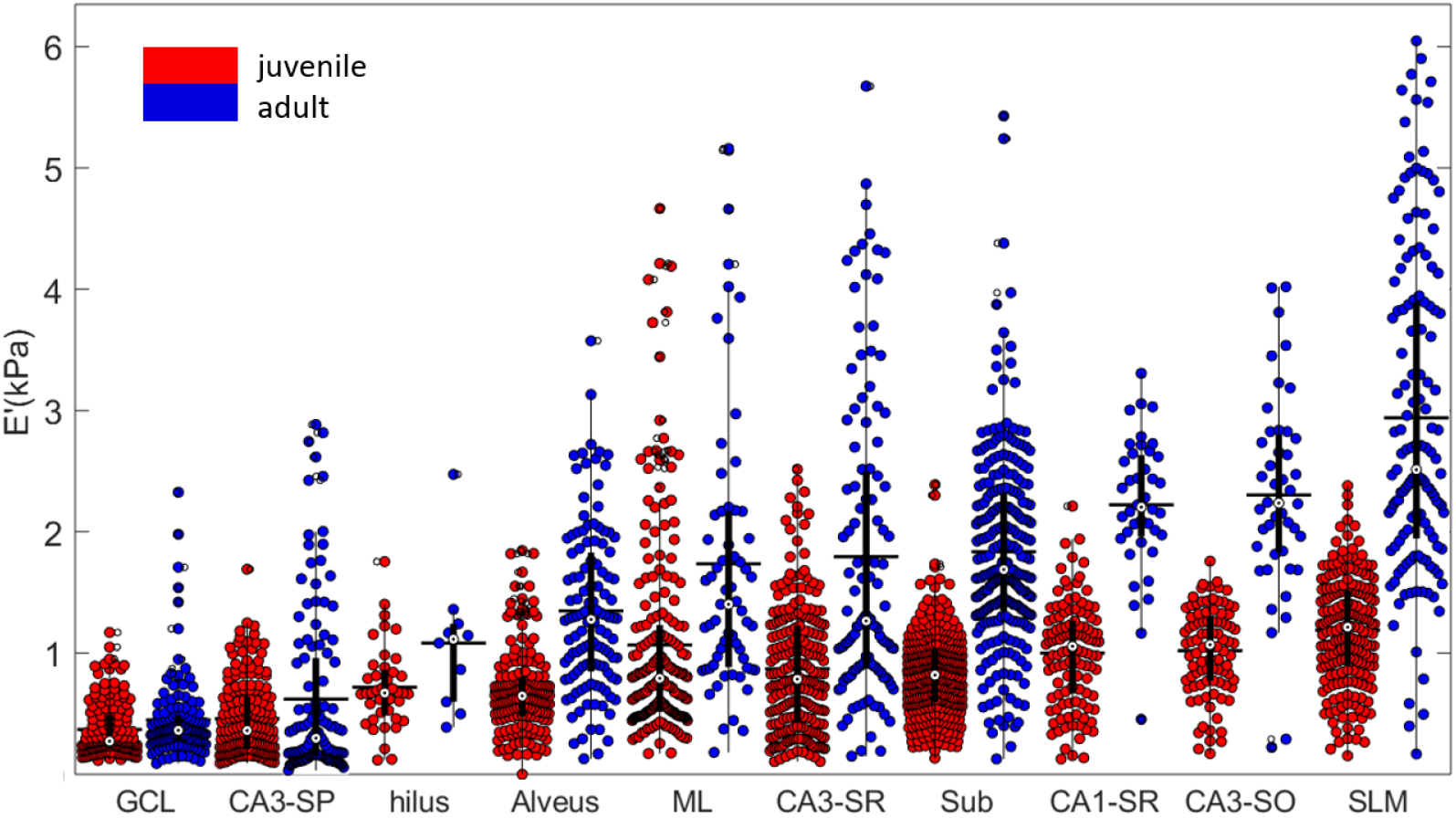
Comparison of storage modulus of juvenile (red) and adult (blue) hippocampal subregions.

**Figure 2:**
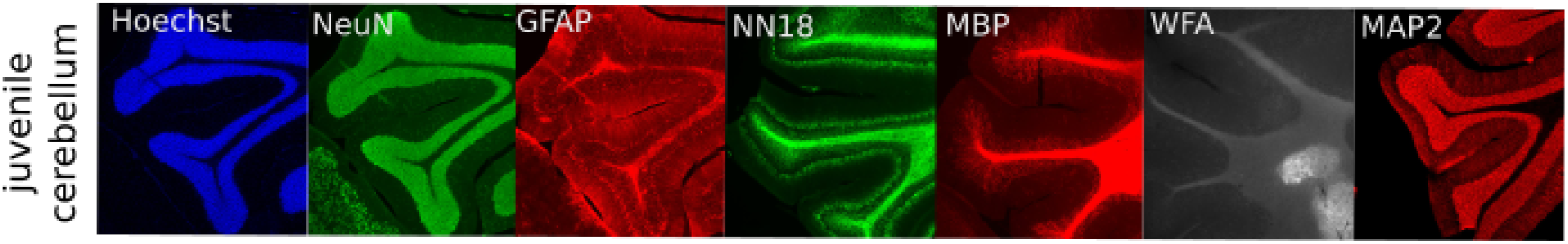
Fluorescent images of immunohistochemical stainings of juvenile mouse cerebellum. Dimensions of images are 2065 μm (width) and 1878 μm (height).

**Figure 3:**
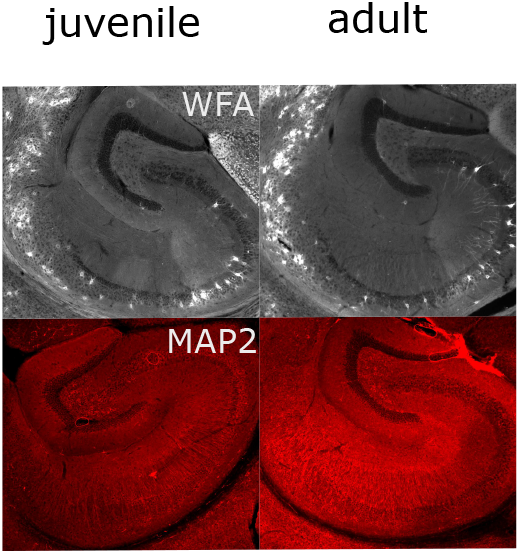
Representative fluorescent images of immunohistochemical stainings of PNNs (WFA) and dendrites (MAP2). Dimensions of images are 2065 μm (width) and 1878 μm (height).

**Figure 4:**
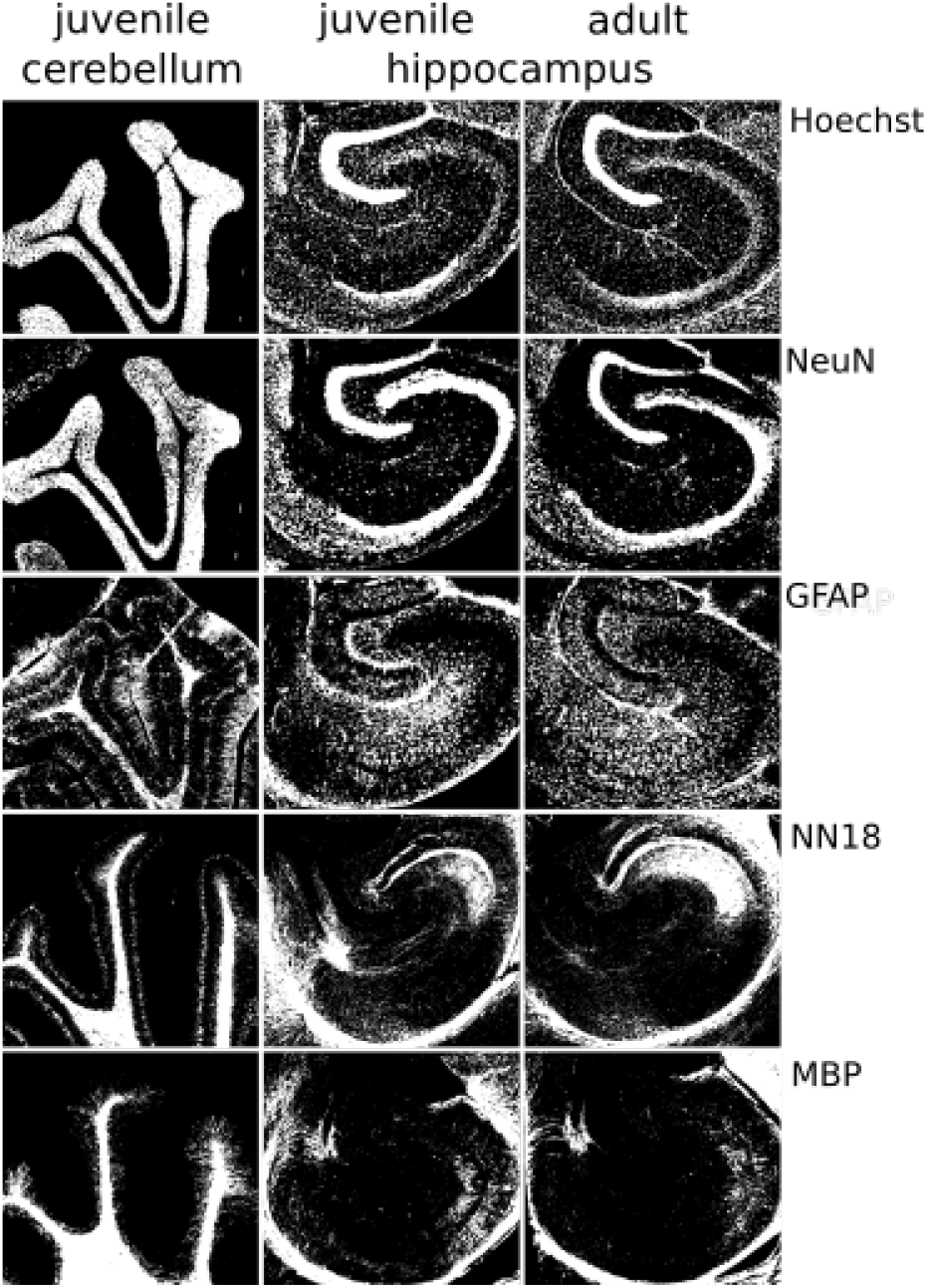
Thresholded fluorescent images used for estimation of relative area covered by stained component. Dimensions of images are 2065 μm (width) and 1878 μm (height).

**Table 3:** Relative area covered with stained component for different brain regions of 1 and 6 month old mice.

|  | 1 month old |  |  |  |  |  |  |  |  |  | 6months old |  |  |  |  |  |  |  |  |  |
| --- | --- | --- | --- | --- | --- | --- | --- | --- | --- | --- | --- | --- | --- | --- | --- | --- | --- | --- | --- | --- |
|  | Hoechst (nuclei) |  | NeuN (neurons) |  | GFAP (astrocytes) |  | NN18 (axons) |  | MBP (myelin) |  | Hoechst (nuclei) |  | NeuN (neurons) |  | GFAP (astrocytes) |  | NN18 (axons) |  | MBP (myelin) |  |
|  | A(%) | SD | A(%) | SD | A(%) | SD | A(%) | SD | A(%) | SD | A(%) | SD | A(%) | SD | A(%) | SD | A(%) | SD | A(%) | SD |
| ML | 12.7 | 1.4 | 2.5 | 1.4 | 6.2 | 4.4 | 2.3 | 1.2 | 1.4 | 0.6 | 10.5 | 2.6 | 1.2 | 0.5 | 22.4 | 6.5 | 1.4 | 1.3 | 2.2 | 1.9 |
| GCL | 95.2 | 3.3 | 91.7 | 2.5 | 5.3 | 4.7 | 0.6 | 1.0 | 0.4 | 0.3 | 95.3 | 2.2 | 87.3 | 4.6 | 5.0 | 1.7 | 0.1 | 0.1 | 1.2 | 1.4 |
| Hilus | 20.4 | 2.9 | 4.3 | 2.2 | 15.1 | 8.2 | 11.3 | 8.0 | 3.3 | 3.2 | 17.3 | 5.3 | 2.3 | 0.8 | 28.2 | 7.7 | 11.5 | 5.5 | 7.7 | 6.2 |
| CA3-SO | 8.4 | 1.8 | 1.8 | 1.3 | 4.6 | 3.1 | 5.4 | 4.4 | 15.3 | 8.5 | 8.0 | 1.6 | 1.8 | 0.6 | 9.1 | 4.2 | 20.5 | 15.0 | 8.2 | 4.7 |
| CA3-SP | 31.7 | 9.8 | 79.7 | 12.8 | 0.2 | 0.1 | 30.9 | 5.7 | 9.4 | 6.1 | 33.1 | 10.6 | 75.4 | 9.6 | 2.0 | 1.3 | 33.5 | 7.3 | 11.7 | 6.4 |
| CA3-SR | 7.0 | 1.0 | 2.0 | 1.1 | 3.2 | 2.0 | 27.9 | 11.6 | 9.4 | 5.3 | 6.4 | 1.4 | 0.8 | 0.3 | 9.2 | 2.4 | 42.6 | 21.0 | 11.9 | 7.7 |
| SLM | 13.0 | 2.0 | 1.0 | 0.4 | 15.9 | 4.0 | 7.0 | 3.2 | 2.7 | 1.5 | 11.5 | 1.9 | 0.6 | 0.2 | 25.1 | 5.2 | 7.4 | 3.0 | 8.2 | 6.7 |
| CA1-SO | 12.0 | 3.0 | 3.0 | 1.6 | 6.9 | 4.0 | 21.6 | 12.0 | 13.1 | 8.0 |  |  |  |  |  |  |  |  |  |  |
| CA1-SP | 51.4 | 11.0 | 63.0 | 15.1 | 4.4 | 3.0 | 4.2 | 2.3 | 13.7 | 9.0 |  |  |  |  |  |  |  |  |  |  |
| CA1-SR | 8.0 | 1.0 | 1.0 | 0.7 | 11.2 | 5.0 | 5.5 | 1.5 | 2.5 | 2.0 | 5.9 | 1.2 | 1.2 | 0.5 | 21.9 | 5.0 | 6.1 | 4.3 | 1.8 | 1.0 |
| Sub | 21.0 | 3.0 | 26.0 | 10.6 | 2.7 | 2.0 | 3.7 | 2.0 | 12.5 | 10.0 | 14.1 | 3.2 | 21.0 | 2.3 | 7.9 | 2.6 | 3.0 | 2.2 | 15.8 | 8.2 |
| Alv | 28.7 | 6.0 | 1.0 | 0.7 | 31.8 | 13.0 | 92.5 | 4.3 | 93.8 | 5.0 | 20.7 | 3.2 | 0.7 | 0.6 | 30.8 | 8.4 | 85.3 | 3.8 | 95.5 | 2.8 |
| Cer-ML | 1.0 | 1.0 | 0.0 | 0.2 | 5.7 | 3.0 | 4.3 | 2.5 | 0.1 | 0.0 |  |  |  |  |  |  |  |  |  |  |
| Cer-GCL | 94.0 | 4.5 | 76.0 | 9.0 | 6.7 | 3.0 | 17.8 | 4.9 | 10.4 | 4.0 |  |  |  |  |  |  |  |  |  |  |
| Cer-Av | 15.0 | 6.0 | 4.0 | 2.0 | 49.4 | 12.0 | 96.4 | 1.1 | 94.7 | 3.0 |  |  |  |  |  |  |  |  |  |  |

**Table 4:**
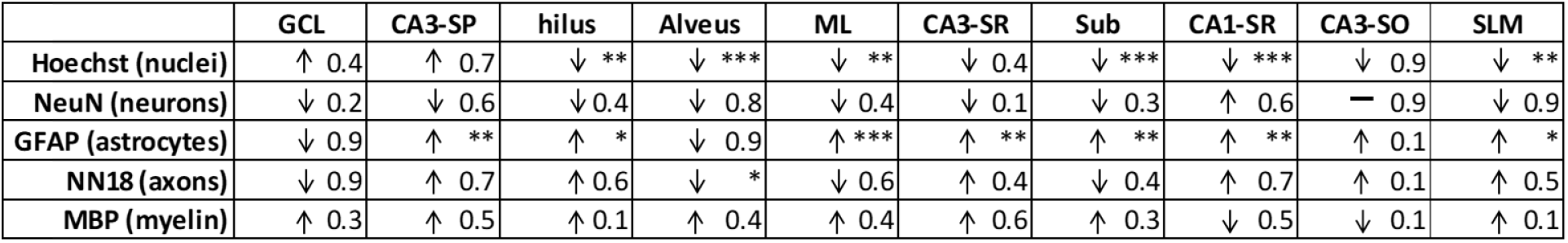
Statistical differences in terms of p-values between juvenile and adult brain components of hippocampal regions. Arrows indicate an increase or decrease in A_*rel*_. ***p<0.001,**p<0.01, *p<0.5, ns - non significant.

